# Unicellular life balances asymmetric allocation and repair of somatic damage representing the origin of r/K selection

**DOI:** 10.1101/2023.11.21.568103

**Authors:** Dmitry A. Biba, Yuri I. Wolf, Eugene V. Koonin, Nash D. Rochman

## Abstract

Over the course of multiple divisions, cells accumulate diverse non-genetic, somatic damage including misfolded and aggregated proteins and cell wall defects. If the rate of damage accumulation exceeds the rate of dilution through cell growth, a dedicated mitigation strategy is required to prevent eventual population collapse. Strategies for somatic damage control can be divided into two categories, asymmetric allocation and repair, which are not, in principle, mutually exclusive. Through mathematical modelling, we identify the optimal strategy, maximizing the total cell number, over a wide range of environmental and physiological conditions. The optimal strategy is primarily determined by extrinsic (damage-independent) mortality and the physiological model for damage accumulation that can be either independent (linear) or increasing (exponential) with respect to the prior accumulated damage. Under the linear regime, the optimal strategy is either exclusively repair or asymmetric allocation whereas under the exponential regime, the optimal strategy is mixed. Repair is preferred when extrinsic mortality is low, whereas at high extrinsic mortality, asymmetric damage allocation becomes the strategy of choice. We hypothesize that optimization over somatic damage repair and asymmetric allocation in early cellular life forms gave rise to the *r* and *K* selection strategies.

## Introduction

Over the course of multiple divisions, cells accumulate two types of damage, genetic and somatic. Somatic damage includes the deterioration of all cellular components other than genomic DNA, including RNA (Macara & Mili, 2008), proteins (Chondrogianni et al., 2014), and other macromolecular structures, such as cell walls (Molina-Quiroz et al., 2013; Vigouroux et al., 2020). Perhaps the most common form of somatic damage is the accumulation of misfolded proteins in the cytoplasm (Coelho et al., 2013, 2014; Dougan et al., 2002; Kopito, 2000; Nakaoka & Wakamoto, 2017; Vedel et al., 2016). Somatic damage, like genetic damage, is heritable across cell divisions, but is not copied prior to its distribution among daughter cells. Also like genetic damage, which, in the absence of dedicated mitigation leads to the eventual collapse of asexual populations through Muller’s ratchet (Govindaraju et al., 2020; Muller, 1964), somatic damage will lead to collapse when damage accumulates faster than it is diluted through cell growth and division.

Somatic damage mitigation strategies can be coarsely divided into two categories: asymmetric allocation and repair which are not, in principle, mutually exclusive. Repair consists of the synthesis of replacement materials for existing essential machinery (e.g. correcting cell wall defects (Vigouroux et al., 2020)); degradation of damaged components within the cell into reusable materials (e.g. proteolytic degradation (Dougan et al., 2002)); or extrusion of dysfunctional material into the external environment (e.g. misfolded protein aggregates (Jang et al., 2010)). All these are active strategies, requiring variable energy investment. The most widespread mechanism of somatic damage repair appears to be proteolytic degradation of misfolded proteins (Dougan et al., 2002). Asymmetric damage allocation is also an active strategy, requiring intracellular shuttling, but does not require investment in synthesis, degradation, or extracellular extrusion. This potential energetic benefit of asymmetric damage allocation comes at a cost: the fitness of one daughter cell, which receives less than half of the somatic damage accumulated by the mother cell, increases at the expense of the other daughter cell which receives more than half.

This tradeoff resulted in the evolution of cell growth strategies including asymmetric allocation of both resources and somatic damage across the tree of life. Morphologically asymmetric cell division, in which the asymmetric allocation of resources and/or damage is apparent after a single cell division, is most widely documented as it can be observed using standard light microscopy during short experiments (Ackermann et al., 2003; Aguilaniu et al., 2003; Sinclair & Guarente, 1997). A thoroughly studied model organism, the bacterium *Caulobacter crescentus*, undergoes morphologically asymmetric division to yield one stationary, stalked, cell and one motile, swarmer cell. The stalked cells live on surfaces in freshwater lakes (Govers & Jacobs-Wagner, 2020) and exhibit senescence, a reduced reproductive output over time (Ackermann et al., 2003). Every pole of each cell originates as the result of septum formation during a prior cell division. At the start of each cycle, the stalked cell inherits a new pole and the old pole that is fixed to the substrate through the stalk. In addition to the asymmetric inheritance of reduced cell wall integrity that is intrinsic to the underlying morphological asymmetry, under environmental stress including heat shock and antibiotic exposure, *C. crescentus* accumulates somatic damage in the form of cytoplasmic protein aggregates. Under moderate stress, these aggregates are distributed symmetrically and repaired by chaperone DnaK and the disaggregase ClpB. When stress intensifies, or when these repair pathways are inhibited through genetic perturbation, the aggregates are shuttled and polarized within the mother cell, resulting in asymmetric inheritance (Schramm et al., 2019).

In the model eukaryote yeast *Saccharomyces cerevisiae*, cell division is achieved through a process of budding whereby one daughter cell is substantially larger (the lineage of this larger progeny is often referred to as the time-independent mother cell) than the other at the time of septum formation(Talia et al., 2009). In contrast to this asymmetric allocation of resources which favors the larger cell, asymmetric allocation of somatic damage favors the smaller cell. In this system, cytoplasmic protein aggregates are completed retained by the larger cell even under standard culture conditions for many generations. Over many cell cycles, the machinery responsible for this asymmetric allocation degrades in the larger cell and aggregates begin to be distributed symmetrically (Aguilaniu et al., 2003). The mechanisms of aggregate shuttling in *S. cerevisiae* have been extensively investigated. Misfolded proteins are directed into dedicated subcellular compartments known as JUNQ (JUNxtanuclear Quality control compartment) and IPOD (Insoluble PrOtein Deposit) attached to the organelles which are retained by the larger progeny cell. Deletion of the chaperone Hsp104 has been shown to impede JUNQ/IPOD localization and partially inhibit asymmetric allocation (Spokoini et al., 2012). Another chaperone, Hsp42, plays a critical regulatory function in asymmetric allocation. Loss of Hsp42 results in symmetric aggregate allocation which, in turn, increases the maximum number of divisions for the larger progeny lineage (Saarikangas & Barral, 2015). Shuttling appears to be achieved through cytoskeletal engagement. Inhibition of myosin motor protein Myo2p and actin polymerization effectors latrunculin A and formin Bni1p has been shown to impede asymmetric allocation through a mechanism of action involving the polarisome macromolecular complex (Liu et al., 2010). In addition to cytoskeletal transport, shuttling can be achieved through ER/Golgi engagement, the primary mechanism observed for transgenic Huntingtin protein aggregates in model disease systems (Song et al., 2014). The NAD-dependent protein deacetylase Sir2 has been demonstrated to influence ER/Golgi-mediated asymmetry (Aguilaniu et al., 2003; J. Song et al., 2014).

The unicellular algae diatom *Ditylum brightwellii* also engages in an asymmetric division strategy whereby one daughter cell is substantially larger than the other. In this system, the average cell cycle duration of the larger progeny cell is shorter than that of the smaller progeny cell, a feature attributed to asymmetric resource allocation (Laney et al., 2012). Whether active asymmetric somatic damage allocation strategies are also employed by this organism, remains an open question.

In multicellular tissues, development and homeostasis is achieved through both limited symmetric division of fully differentiated cells and asymmetric division of stem cells resulting in one daughter cell that maintains stemness and another that begins to differentiate. During embryogenesis, in *Drosophila melanogaster*, ganglion mother cells (GMCs) divide once to produce two fully differentiated neurons or glial cells. GMCs are themselves the result of an asymmetric division of a precursor neuroblast which produces one neuroblast and one GMC. During this division event, the neuroblast, which will undergo apoptosis in a later stage of embryogenesis, retains all protein aggregates ensuring that the fully differentiated tissue is free from somatic damage (Bufalino et al., 2013; Rujano et al., 2006). Similarly, during homeostasis in both *D. melanogaster* (Bufalino et al., 2013) and human intestinal (Rujano et al., 2006) stem cell crypts, the asymmetric division of intestinal stem cells produces one cell which remains in the stem cell crypt and one cell which begins differentiation. In this case, however, protein aggregates are completely shuttled into the differentiating cell. In both contexts, somatic damage is asymmetrically allocated to the short-lived lineage.

Technological developments in microfluidics (Andersson & van den Berg, 2003) and time-lapse microscopy (Balaban et al., 2004), which enabled tracking many lineages of cells over many cell cycles, revealed that even cells which divide morphologically symmetrically, can implement an asymmetric damage allocation strategy. Similarly to *C. crescentus* (see above), in the morphologically symmetric bacteria *Escherichia coli*, symmetry breaking between the poles of the cell was observed. Every pole originates as the result of septum formation during a prior cell division. At the start of each cycle, the cell inherits one new pole and one old pole of variable age. Cells that inherit older poles show increased death rates and decreased growth rates compared to cells inheriting younger poles (Stewart et al., 2005). As in the morphologically asymmetric organisms described above, this effect is primarily attributed to the asymmetric accumulation of protein aggregates (Lindner et al., 2008).

Similar dynamics have been directly observed in the morphologically symmetric bacteria, and intracellular pathogen, *Mycobacterium tuberculosis*. In this system, irreversibly oxidized proteins (IOPs) accumulate under host-induced stress. IOPs are not repaired but asymmetrically allocated between daughter cells at division. Daughter cells which do not receive IOPs have a shorter cell cycle duration on average and reduced mortality following antibiotic exposure. Chaperone ClpB has been shown to facilitate asymmetric allocation whereas the absence of ClpB is associated with reduced fitness in response to environmental stress *in-vitro* and reduced virulence *in-vivo* (Vaubourgeix et al., 2015).

The symmetric model eukaryote, fission yeast *Schizosaccharomyces pombe*, also asymmetrically allocates protein aggregates resulting from both heat and oxidative stress (Coelho et al., 2013). Although the senescence of this organism remains a matter of debate (Nakaoka & Wakamoto, 2017), asymmetric allocation has been demonstrated to be an active process mediated by the Hsp16 chaperone (Coelho et al., 2014).

Asymmetric allocation in morphologically symmetric species can also be environment-dependent. In *S. pombe*, symmetry breaking is only observed under stress(Coelho et al., 2013, 2014; Nakaoka & Wakamoto, 2017). In *E. coli*, asymmetric allocation of protein aggregates also becomes more pronounced in stress conditions, for example, under heat shock (Vedel et al., 2016). These findings highlight the phenotypic plasticity of asymmetric allocation as a damage control strategy.

While it is apparent that mechanisms for both repair and asymmetric allocation of somatic damage have evolved across the tree of life, the environmental and physiological determinants of optimal strategy remain poorly understood. Several mathematical modeling approaches have been applied to this problem(Blitvić & Fernandez, 2020)(Borgqvist et al., 2020; Schnitzer et al., 2020; R. Song & Acar, 2019)(Min & Amir, 2021)(Watve et al., 2006), the most common one being agent-based simulations (Ackermann et al., 2007; Chao, 2010; Clegg et al., 2014; Koleva & Hellweger, 2015; Wright et al., 2020). The principal advantage of the agent-based approach is the ability to compare relative fitness of agents with selected parameter sets in direct competition, without constructing of an absolute fitness function. Charting the broader landscape of the dependence of the somatic damage mitigation strategy on total population size or growth rate, however, remains challenging.

We sought to construct a framework to assess the impact of both cell physiology and environmental parameters with respect to an absolute fitness function, namely, the total number of clonal cells. We propose a model of somatic damage accumulation among cells in a chemostat, a stable environment which admits an equilibrium total population size limited by nutrient availability, extrinsic mortality, and somatic damage accumulation. We identify the optimal mitigation strategy, maximizing the total population size and explore an expansive domain of the parameter space with more than 2.6 million initial conditions where we assume repair, but not asymmetry, to be explicitly growth costly. Within this framework, we find that the optimal strategy is primarily determined by the extrinsic mortality and the physiological model for damage accumulation, consistent with the interpretation that asymmetric allocation and repair in unicellular life represent, respectively, r and K selection regimes. We hypothesize that, at early stages of cellular evolution, optimization over somatic damage repair and asymmetric allocation played a critical role in establishing repair and asymmetry specialists including the evolution of morphologically asymmetric division. Thus, the balance of repair and asymmetry in early cellular life appears to represent the prototypical r/K selection strategies choice that is likely to be an inalienable feature of the evolution of life.

## Methods

### Model Construction

We begin with the description of the chemostat and an idealized cell population in which the somatic damage concentration rate is the same for all individuals at any time. The chemostat, of volume *V*, contains cell media with dissolved nutrients, concentration *ϕ*(*t* = 0) = *ϕ*_0_. Volume *V*^*e*^ is removed per unit time and replaced with fresh cell media (concentration *ϕ*_0_, increasing the nutrient concentration at rate 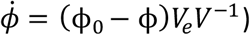. Cell mass is measured in units of nutrient. Without investment in repair (asymmetry is assumed not to be growth costly), *v* volume of media is ingested per unit cell mass, per unit time and 100% of the available nutrient in that volume is converted to cell mass. Investment in repair, *R*, decreases this fraction proportionately. Cell mass density, *x*^−1^, is invariant. So defined, the growth rate for the total cell volume in the chemostat is 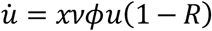 and the removal of nutrient from the environment reduces the nutrient concentration at rate 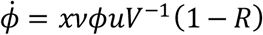.

While cell growth can be defined with respect to a unit cell volume, cell loss is defined with respect to individual cells. However, although the average volume per individual cell can be a time-dependent quantity, when somatic damage concentration is the same for all individuals, the probability of cell loss is independent of individual cell volume. Thus, it is possible to describe the rate of the total cell volume loss, without loss of generality.

Cells within volume *V*^*e*^ are removed from the chemostat, representing extrinsic mortality that by definition does not depend on somatic damage accumulation. Intrinsic mortality is determined by accumulated somatic damage. The concentration of somatic damage, ρ, increases at a constant rate 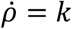 independent of prior accumulated damage or at a variable rate 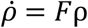, representing linear and exponential damage accumulation, respectively. Across all conditions studied, *k*>0 but *F* ≥ 0. Somatic damage concentration also decreases due to cell growth and repair at a rate 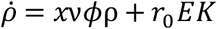 where *E* is a unitless constant of proportionality. The death rate due to accumulated somatic damage is modeled according as 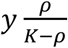, where *K* is the lethal threshold for damage concentration, and [*y*] = *t* ^−1^ is a constant of proportionality. So defined, the total cell volume in the chemostat decreases at the rate 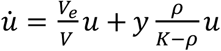.

We can now proceed with the nondimensionalization of this system. We are free to set the timescale with respect to the constant of proportionality, *y* ≡ 1. The units of nutrient and damage concentration are normalized with respect to ϕ_0_ and *K*; for convenience, we introduce 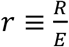. This yields a complete model description, the system of ordinary differential equations (ODEs):

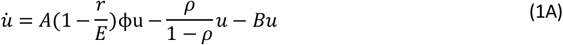

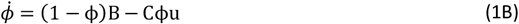

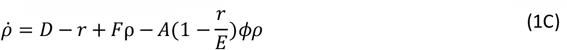

where *A* ≡ *xv*ϕ_0_, representing growth; 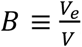, representing dilution; 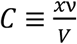, representing nutrient acquisition; 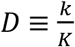, representing linear damage accumulation; *F*, representing exponential damage accumulation; and *r*, representing repair, are rate parameters, [:] = *t* ^−1^. The final parameter, *E*, is unitless, and represents repair efficiency. When *E* → ∞, a negligible fraction of the nutrient acquisition is required to repair all somatic damage. There exists a unique, non-trivial (*n*^∗^ > 0), equilibrium solution, when the growth rate is suitably large (see Appendix I). In the limit 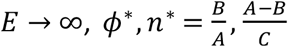 and the optimal damage mitigation strategy maximizes investment in repair, *r* = *D*. In the limit *E* → 0, the optimal strategy is not to invest in repair, *r* = 0. More generally, this framework supports evaluating the optimal investment in repair, but not evaluating the impact of asymmetric damage allocation, because it assumes that somatic damage concentration is the same for all individuals at any time.

Asymmetric allocation is modeled by Introducing the parameter *a* determining the fraction of damage inherited by each daughter cell,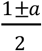 . Over many generations, and therefore at equilibrium, asymmetric allocation results in a continuum of possible damage concentrations for different cell lineages. Under the simplifying constraint that the probability of cell division is independent of the time elapsed since the previous cell division, the partial differential equation (PDE) below describes the chemostat as introduced above (see Appendix II for derivation):

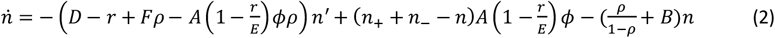

where *n*(*ρ, t*) is the number of cells with damage concentration 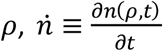 represents the time derivative as above, 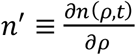 represents the derivative with respect to damage concentration, and 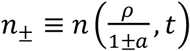. The dependence on the shifted terms *n*^±1;^ make this system poorly suited for investigation using analytical approaches that are applicable for most first-order PDEs; however, recently techniques have been developed for equations of a similar form describing related biological systems (Pikovsky & Tsimring, 2023).

Relieving the constraint that the probability of cell division is independent of the previous cell division, requires explicitly modelling an individual cell cycle, through recording an additional state variable. We implemented an idealized cell cycle in which cell division occurs deterministically when the cell reaches a critical mass 2*p*_0_ and divides morphologically symmetrically so that each daughter cell inherits a mass of *p*_0_. This yields the system:

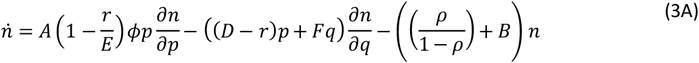

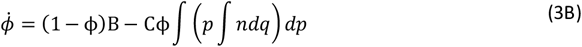

where *p* is the amount of accumulated nutrient and *F* = *ρp* is the amount of accumulated damage (see Appendix III for derivation). Within this formalism, asymmetric damage allocation is introduced through the equilibrium boundary conditions:

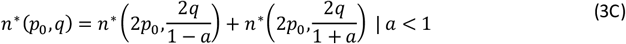

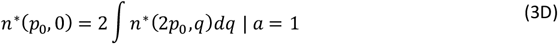

This system can be solved using the method of characteristics for appropriate boundary conditions. The solutions for the special cases *a* = 0,1 are described in Appendix IV.

For any value of *a*, there exists at most one nontrivial equilibrium solution (see Appendix V for proof); however, even for the special cases, *a* = 0,1, computing the solution requires several numerical steps. Furthermore, the optimal strategy for *F* > 0 can, in principle, include high rates of repair, *r* > *D*; however, in this formalism, *r* > *D* is a non-physical solution when *q* is small. For these reasons we interrogated this system with a fully discrete description of *p* and *q* according to the following stochastic master equation:

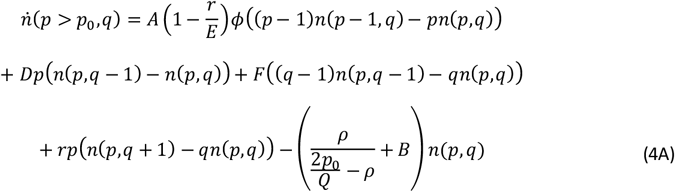

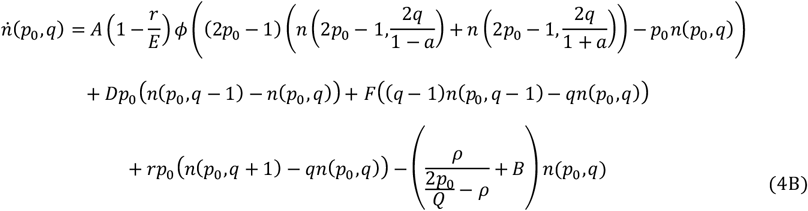

dependent on *p*, where cell division (from one cell in state 2*p*_0_ to two cells in state *p*_0_) is deterministic, reflected by the direct transition from cells in state 2*p*_0_ − 1 to state *p*_0_. The two parameters that define the discretization of the system are *p*_0_, which determines the number of reactions required to complete an entire cell cycle and *Q* which determines the maximum discrete units of somatic damage which may be accumulated by any cell. 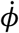 is expressed as in eq. 3B with the integrals replaced by sums.

### Exploration of the Parameter Space

The equilibrium solution depends on both discretization parameters such that *p*_0_ determines the number of reactions required to complete an entire cell cycle and *Q* determines the ratio of the number of reactions in which damage is accumulated to that in which damage is repaired which results in cell death. Although the model dependence is nonlinear with respect to both *p* and *q*, all possible values of *p* are within a factor of two, *p*_0_ ≤ *p* < 2*p*_0_ whereas the ratio between the maximum and minimum values of *q* is unbounded, 0 ≤ *F* < *Q*. Thus, the solution is more sensitive to *Q* than to *p*_0_. In all simulations, *p*_0_, *Q* = 40,1000 were fixed to address this imbalance and achieve stability with respect to discretization so that the behavior approaches the limit *p*_0_, *Q* → ∞ (see Fig. S1).

The parameters *A* and *C* could be fixed as well. The optimal strategy was assessed for environments supporting cell populations large enough for stochastic fluctuations around the average population size not to result in extinction. It follows that the ratio of parameters 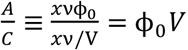, representing the maximum amount of nutrient available in the chemostat at any time, is arbitrary with respect to the determination of optimal strategy whenever *A* ≫ *C* . Jointly decreasing *A, C* corresponds to increasing the cell cycle duration for a given environmental condition by either lowering the nutrient density, ϕ_0_ ≪ 1, *V* ≫ 1 or lowering the rate at which cells extract nutrient from the environment, *xv*. Increasing the cell cycle duration impacts the total population size by increasing the probability of cell death per cell cycle; however, it minimally impacts the relative probability of extrinsic mortality to intrinsic mortality (as the result of accumulated somatic damage). We validated through simulation that for fixed *A*, varying *C* impacted the total population size but not the optimal strategy at equilibrium (see Fig. S2). We proceeded to fix *A* so that unit time is equal to a single cell cycle when *ϕ* = 1, *r* = 0 and *C* = 10^−6^ to be suitably small so that the total population size is on the order of 10^5^ cells for most conditions studied.

For fixed *p*_0_, *Q, A*, and *C*, the parameter ranges *B* ∈ (0, 0.66),*D* ∈ (0, 0.25),*E* ∈ (0, 0.3),*F* ∈ (0, 3) were identified to fully describe all variation in optimal strategy: increasing beyond these bounds either results in extinction or increases the total cell population but does not impact the dependence on the remaining parameters with respect to optimal strategy. For every (*B, D, E, F*) parameter set, the nontrivial solution for eqs. 4 was computed as a function of strategy, *n*^∗^(*a,r*) and the optimal strategy, maximizing 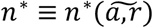, was identified (see Table 1 for a summary of parameters and Appendix VI for details regarding the numerical methods).

**Table 1.** Parameters used in simulations. The last column gives measurements of parameters in different natural systems, both directly and indirectly related to the parameters used in the model.

| Parameter | Range | Description | Related Measurements<sup>a</sup> |
| --- | --- | --- | --- |
| $A$ | 28.0 | Growth rate | Cell cycle duration in *S. pombe* on agarose at 30°C: 133 minutes (Coelho et al., 2013); cell cycle duration in *E. coli* on agarose at 37°C: 25 minutes (Vedel et al., 2016). |
| $B$ | 0-0.66 | Dilution rate | Predatory mortality in planktonic bacteria: 1-10%, 30-45% (Pace, 1988). |
| $C$ | 1e-6 | Nutrient acquisition rate | Uptake rate of glucose by Baltic Sea beach bacteria: $4.7 * 10^3 \frac{\mu g}{g * h}$ (Meyer-Reil, 1978). |
| $D$ | 0-0.25 | Linear component of damage accumulation | Rate of aggregate formation in *S. pombe*: 14 aggregates/cell cycle (Coelho et al., 2014); rate of aggregate formation in *E. coli* under heat stress: $0.03 \mu m^2$/cell cycle (Vedel et al., 2016). |
| $E$ | 0-0.3 | Repair efficiency | Cost of refolding by Hsp70 for 564 amino acids long protein: 5 ATP molecules (Fauvet et al., 2021). |
| $F$ | 0-3 | Exponential component of damage accumulation | Aging rate accelerates with age in *C. crescentus*: $F \gg 0$ (Ackermann et al., 2003). |
| $a$ | 0-1 | Level of asymmetry | Asymmetry in protein aggregates segregation in *S. pombe*: 0 at 30°C – 0.8 at 40°C (Coelho et al., 2013). |
| $r$ | 0-max(D, E) | Rate of repair | Aggregate refolding rate by $3.5 \mu M$ DnaK, $0.7 \mu M$ DnaJ and $3.5 \mu M$ GrpE *E. coli* chaperones: $12 \frac{\mu M}{min}$ (Diamant et al., 2000). |
| $p_0$ | 40 | Amount of nutrient required for division | Birth volume of *S. pombe*: $70 \mu m^3$ (Vyas et al., 2021); birth volume of *E. coli*: $0.6 \mu m^3$ (Kubitschek, 1990). |
| $Q$ | 1000 | Lethal amount of damage at division volume. | *S. pombe* lineages that accumulate more than 5 protein aggregates ultimately die (Coelho et al., 2013). |

### Comparing fitness under different models of damage accumulation

It is useful to identify (*B, D, E, F*) parameter sets for which the optimal fitness, 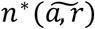, are the same. Consider the parameter set *S*^*F*^ ≡ (*B, D*^*F*^, *E, F*). We may identify the corresponding *D*^0^ > *D*^*F*^ defining the parameter set *S*^0^ ≡ (*B, D*^0^, *E*, 0) such that ^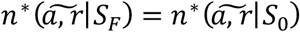^. Comparing the optimal strategy between *S*_0_ and *S*_1_ reveals the relative impact of asymmetry and repair for different models of somatic damage accumulation when the optimal fitness is matched. It is also useful to define a measure of equivalency among parameter sets with respect to the naïve fitness, *n*^∗^(0,0). Among parameter sets for which *n*^∗^(0,0) = 0, it is useful to consider the measure:

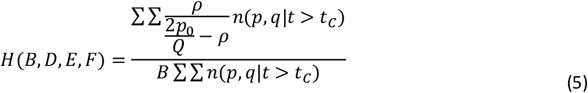

the ratio of the intrinsic and extrinsic mortality rates, which we label “harshness”. This ratio converges by time *t*^*C*^, at a rate faster than the total population size converges (Figure S3). It follows that the harshness of a parameter set may be computed even when *n*^∗^(0,0) = 0. Comparing 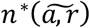 between parameter sets with equal harshness reveals the relative impact of asymmetry and repair for different models of somatic damage accumulation when the initial conditions are matched. In particular, evaluating harshness may reveal equivalent initial conditions for which only select models of somatic damage accumulation admit 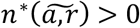.

For an individual parameter set (*B, D, E, F*), it is also useful to have a measure for the relative impact on somatic damage accumulation with respect to *D* and *F*:

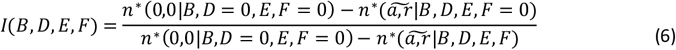

the ratio of the fitness reduction resulting from only the linear damage accumulation term, *D*, to that of both *D* and *F*.

## Results

### Three Optimal Strategies of Somatic Damage Control

Across all conditions studied, all local optima can be classified into: i) asymmetry preferred (*a* = 1,*r* = 0), ii) repair preferred *a* = 0, *r* = *r*_*opt*_, and iii) mixed strategy *a* = 1, *r* = *r*_*opt*_. All optima are sensitive to perturbation with respect to both *a* and *r* but repair preferred optima are substantially less sensitive to perturbation in *a*. Although no local optima were observed for intermediate asymmetry, 0 < *a* < 1, *n*^∗^(*a,r*) can be nonmonotonic with respect to *a*. Thus, optimality of intermediate asymmetry depends on the functional form of the death rate due to somatic damage accumulation, 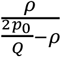, as well as the assumption that asymmetry is not explicitly growth costly (see Discussion). All three types of local optima can be represented within the *n*^∗^(*a,r*) landscape for a single parameter combination (*B, D, E, F*); however, the most common landscapes observed consist of a single optimum. Landscapes with both a repair preferred and an asymmetry preferred local optima are also common (see Fig. 1).

**Figure 1.**
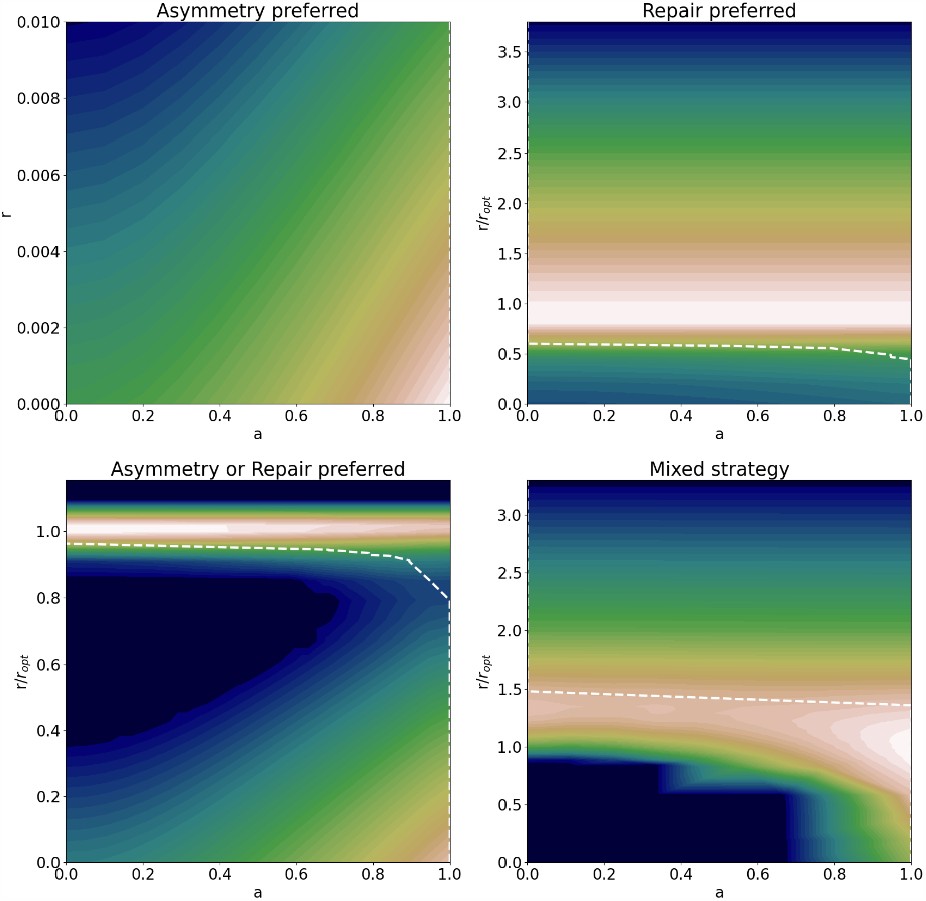
The four most frequently observed fitness landscapes *n*^∗^(*a, r*) for fixed parameters (*B, D, E, F*). The color scheme reflects fitness of *n*^∗^(*a, r*) with blue being the lowest and white being the highest. The dashed line shows the optimal value of asymmetry for a given value of repair. Repair is reported with respect to the optimal value, *r/r*_*opt*_ when *r*_*opt*_ > 0 for at least one local optimum. (asymmetry preferred) B=0.02, D= 0.07, E=0.04 (repair preferred) B=0.04, D=0.013, E=0.141, F=0 (asymmetry or repair preferred) B=0.04, D=0.1184, E=0.141, F=0 (mixed strategy) B=0.62, D=0.003, E=0.118, F=1

Examining the population structure at each of the three possible local optima provides more detailed insight into the fitness effects of asymmetric damage allocation and repair (Fig. 2). A naive population has a broad diagonal structure in which cells inherit, on average, the same amount of damage as they will accumulate over the course of one cell cycle. It follows that dividing cells contain approximately twice the amount of damage as newborn cells. An asymmetry preferred strategy subdivides the population into several parallel trajectories. Under exponential damage accumulation, *I* ≪ 1, the survival probability of cells that inherit any amount of damage may be low and only the first trajectory may be substantially represented. In contrast, a repair preferred strategy homogenizes the damage distribution with respect to cell size. This observation suggests repair preferred strategies may be observed more frequently under conditions where *I* ≪ 1 (see below). A mixed strategy combines the effects of subdivision and homogenization, generating a subpopulation with a lower probability of death due to damage accumulation than in a symmetric population with the same investment in repair. The associated cost is the generation of subpopulations with a higher probability of death.

**Figure 2.**
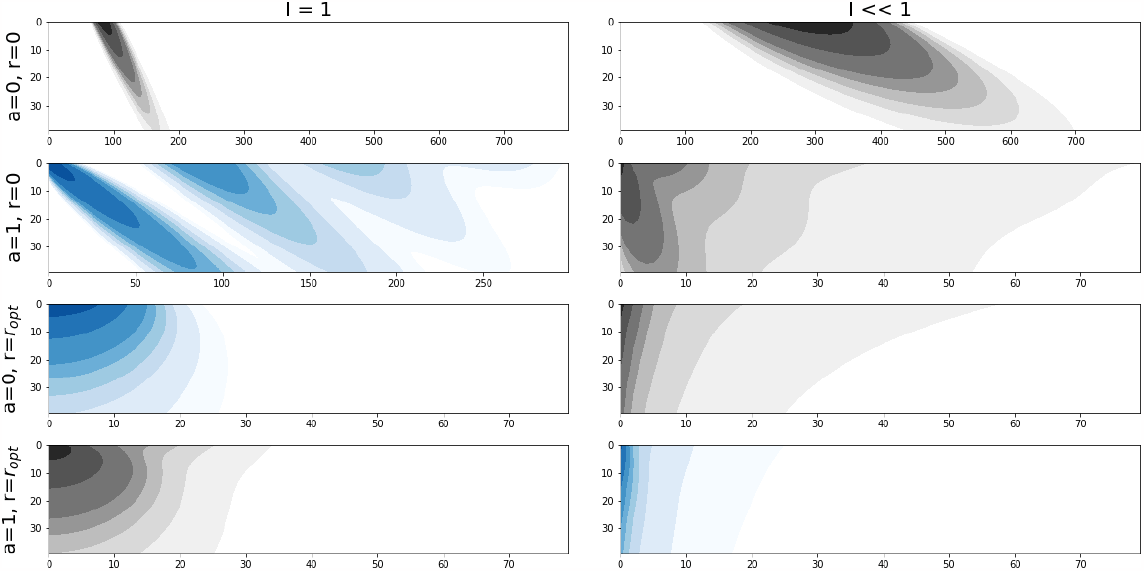
Contour plots of population structures. On each panel x-axis represents damage, *Q*, and y-axis represents amount of nutrient, *p*. The left column corresponds to the asymmetry or repair preferred landscape in Fig. 1 (B=0.4, D=0.053, E=0.176, F=0). The right column corresponds to the mixed strategy landscape (B=0.62, D=0.003, E=0.118, F=1). Local optima are highlighted in blue and structures of related sub-optimal strategies are shown in gray. Shaded regions encompass to 10, 30, 50, 60, 70, 80, 90 and 95%% of the population expanding from high to low population density (shading from dark to light).

From this observation it follows that *n*^∗^(*a,r*) may be non-monotonic with respect to *a* and that *a* is either maximized or minimized for all local optima where *r* > 0. The optimal investment in repair for an individual cell, maximizing its expected number of progeny, depends on the damage it inherits. Increasing *a* alters the inherited damage distribution so that for fixed *r*, cells inheriting the least damage become increasingly invested in repair whereas those inheriting the most damage become increasingly less invested in repair. The optimal value for *a* balances these, nonlinear, effects. When *r* = *r*_*opt*_ the optimal value of *a* is either 0 or 1. When *a* = 0, the investment in repair is optimized. When *a* = 1, the population maximizes its utilization of asymmetry, which is not explicitly growth costly, at the cost of misallocating investment in repair. As with the repair preferred strategy, this observation suggests that the mixed strategy *r* = *r*_*opt*_, *a* = 1 would be observedmore frequently under conditions where *I* ≪ 1 and cells that inherit more damage also accumulate damage more quickly.

### Determinants of Optimal Strategy Represent r/K Selection

After analyzing the range of possible local optima, we proceeded to establish the environmental and physiological determinants of global optimal strategy, 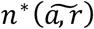, parameterized by (*B, D, E, F*). A map indicating the regions of the parameter space occupied by each possible strategy (repair preferred, asymmetry preferred, and mixed) is shown in Fig. 3. For fixed (*E, F*), 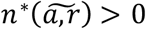 within a triangle bounded by increasing extrinsic mortality (*B*, x-axis) and increasing damage accumulation (*D*, y-axis). Increasing repair efficiency, *E*, increases both the size of this triangle and the fraction occupied by the repair preferred strategy. Within each triangle, the repair preferred region covers an area of reduced damage accumulation and extrinsic mortality with respect to the asymmetry preferred region. Increasing *F* decreases the size of the triangle, as well as the fraction occupied by the asymmetry preferred strategy, and increases the fraction occupied by the mixed strategy.

**Figure 3.**
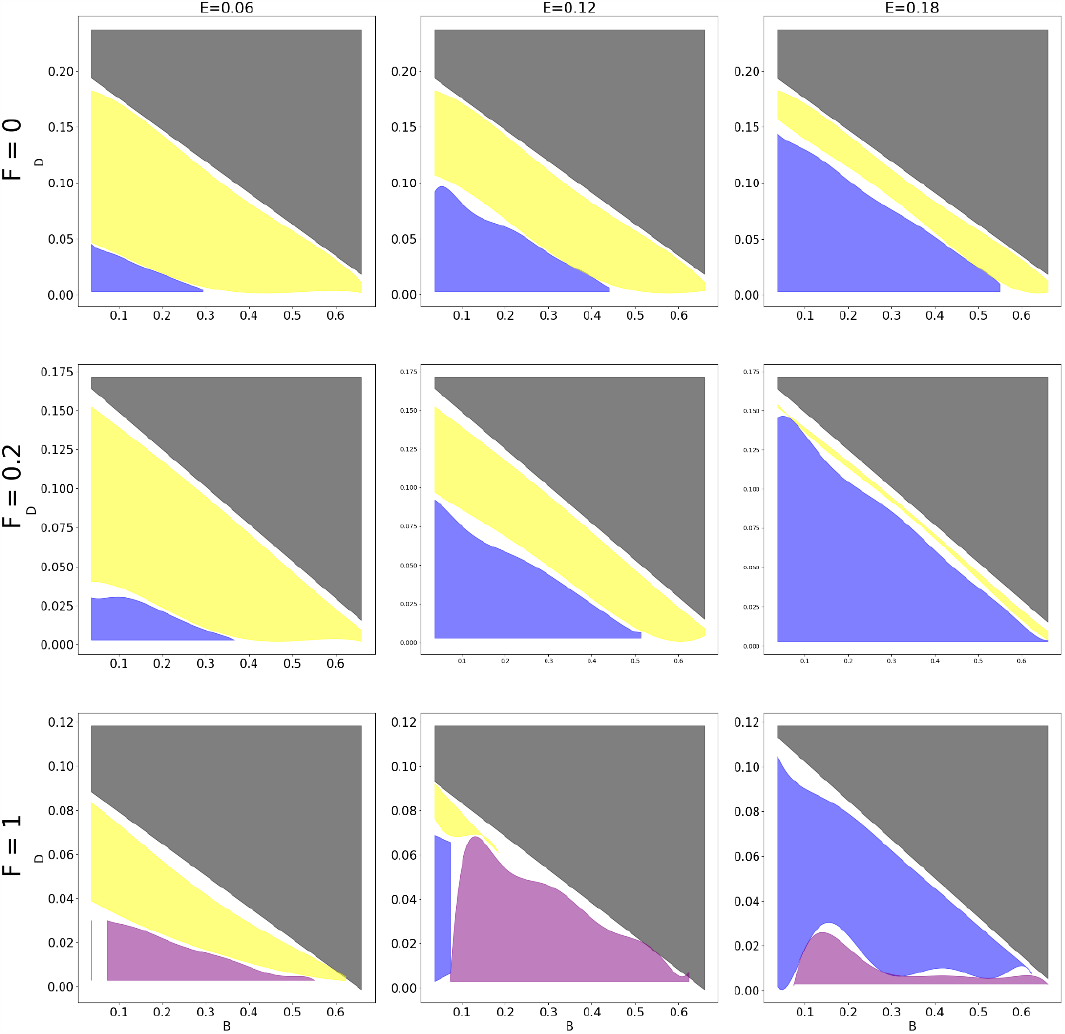
The three strategies of somatic damage control. The map shows which of the three possible strategies (repair preferred, blue; asymmetry preferred, yellow; and mixed, purple) is optimal dependent on (*B, D, E, F*). Within the gray region, 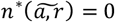. Globally, higher extrinsic mortality, increased *B*, corresponds to an increased prevalence of the asymmetry preferred strategy.

While intrinsic mortality depends on multiple parameters, extrinsic mortality is specified explicitly by *B*. Globally, increasing *B* increases the fraction of the region where 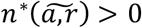occupied by the asymmetry preferred strategy. Investment in repair, which is growth costly, increases the probability of cell death due to extrinsic factors per cell cycle. This trend represents a form of r/K selection, the theory in evolutionary ecology that centers on the trade-off between offspring number and quality (Macarthur & Wilson, 1967; Tenkeu et al., 2020; Theodosius Dobzhansky, 1950). The repair preferred strategy is a K-strategy, producing high-quality offspring, with low rates of intrinsic mortality, at a low growth rate. The asymmetry preferred strategy is an r-strategy, producing offspring of variable quality, some with high rates of intrinsic mortality, at a high growth rate. Typically of r/K selection (Tenkeu et al., 2020), the asymmetry preferred r-strategy is more commonly observed when the extrinsic mortality rate is high, and intrinsic mortality has a reduced impact on the probability of cell death during one cell cycle.

### Optimal Strategy Bifurcates Depending on the Model of Damage Accumulation

Unlike extrinsic mortality, intrinsic mortality depends on multiple parameters, including strategy parameters *a* and *r*, and thus, the dependence of the optimal strategy on intrinsic mortality is complex. We began assessing this dependence at the edge of the region where 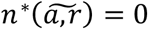, that is, the extinction boundary. First, we established the harshness, *H*, of the conditions parametrized by (*B, D, E, F*) which lie along the extinction boundary. As described under Methods, *H* measures the impact of intrinsic mortality on a naïve population, *n*^∗^(0,0). Increasing *F*, the exponential component of somatic damage accumulation, increases *H* at the extinction boundary (Fig. 4a). It follows that for equally challenging initial conditions, as reflected by equal values of *H*, a mixed somatic damage mitigation strategy incorporating both repair and asymmetry is more likely to prevent population collapse when somatic damage accumulates nonlinearly. This effect can be understood as a consequence of the fact that under a linear regime, the rate of damage accumulation does not depend on the mitigation strategy but under a nonlinear regime, this rate is reduced through mitigation.

**Figure 4.**
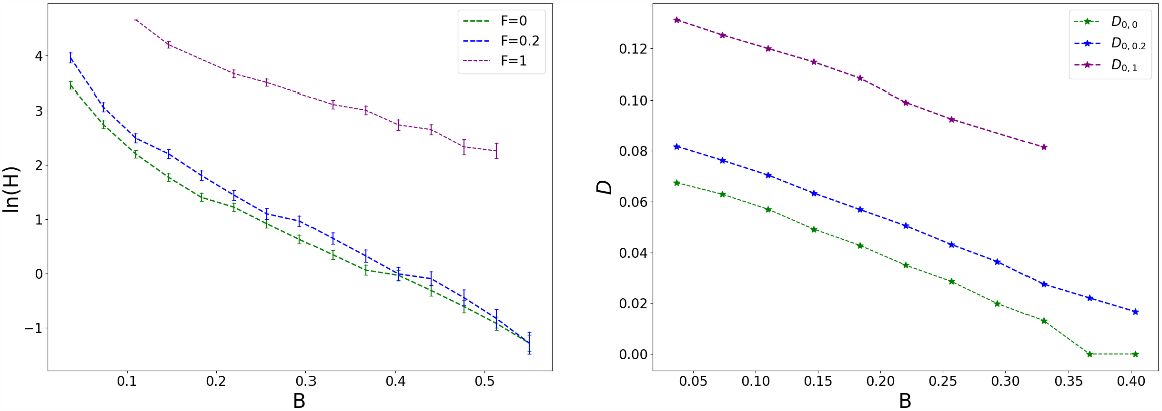
Choice between somatic damage mitigation strategies depending on the model of damage accumulation. (a) Harshness, *H*, at the extinction boundary is higher for increasing *F*. Intervals indicate the largest value sampled where 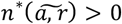 and the smallest where 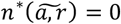. (b) The transition to the asymmetry preferred strategy occurs at smaller 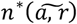 for larger *F* (*D*_0,*F*1_ > *D*_0,*F*2_ > *D*_0,0_ when *F*1 > *F*2). (a&b) *E* = 0.1 for all data shown. Note *H* and therefore *I* vary over each curve.

We proceeded to establish the manner in which the optimal strategy depends on the model of somatic damage accumulation within the region where 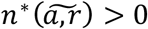. Similar to the analysis described above, we considered conditions at the boundary between the repair preferred and asymmetry preferred regions. For fixed (*B, E, F*), increasing *D* always crosses this boundary with the same orientation, from repair to asymmetry, whenever both regions are present. As described in the Methods, for any parameter set *S*^*F*^ ≡ (*B, D*^*F*^, *E, F*), the corresponding *D*^0,*F*^ > *D*^*F*^ can be identified defining the parameter set *S*^0,*F*^ ≡ (*B, D*^0^,*E*, 0) such that 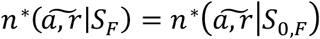. We identified such sets, *S*^*F*^, that lie on the boundary between the asymmetry and repair preferred regions and compared *D*^0,*F*^ to *D*^0,0^, corresponding to where this boundary is crossed when *F* = 0. We observed *D*^0,*F*^ > *D*^0,0^ with the strength of the inequality increasing with the increase of *F* (Fig. 4b). This is equivalent to the statement that the transition to the asymmetry preferred strategy occurs at smaller 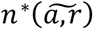 for larger *F*. It follows that, for equally challenging equilibrium conditions, as reflected by equal values of 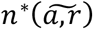, the repair preferred strategy is more commonly observed when damage accumulation is nonlinear and such that the damage accumulation rate is more readily modifiable through investment in repair.

The factors defining the boundary of the region occupied by the mixed strategy are more complex. The prevalence of the mixed strategy is non-monotonic with respect to *E*, and for fixed (*E, F*) the region occupied by the mixed strategy can be adjacent to either only asymmetry preferred, only repair preferred, or both (see Fig. 3). Clearly, however, the mixed strategy only emerges when *F* > 0. As described above, the relative impact on somatic damage accumulation with respect to *D* and *F* can be assessed via *I*, the ratio of the fitness reduction resulting from *D* alone, to that of both *D* and *F*. Projecting the map shown in Fig. 3 to the *BI* coordinate plane reveals an orderly global trend. When *I* ≪ 1, and damage accumulation is highly nonlinear, the mixed strategy is globally optimal. As *I* increases so that the rate of damage accumulation linearizes, the optimal strategy bifurcates into either repair or asymmetry preferred depending on the extrinsic mortality rate (Fig. 5).

**Figure 5.**
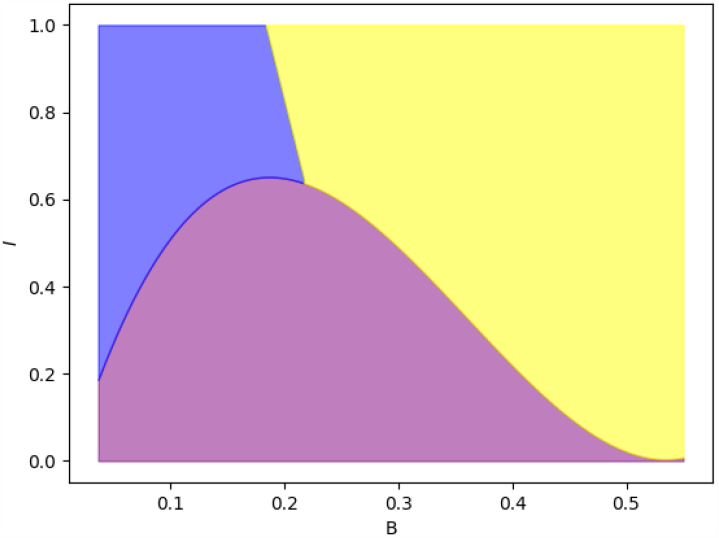
Dependence of the optimal strategy of damage mitigation on the model of damage accumulation. The optimal strategy is mixed (purple) when *I* ≪ 1 such that damage accumulation is highly nonlinear. As *I* increases, and the rate of damage accumulation linearizes, the optimal strategy bifurcates into either repair (blue) or asymmetry preferred (yellow) dependent on whether the extrinsic mortality rate, *B*, is low, or high. Borders between regions are based on the raw data presented on Figure S5.

## Discussion

Somatic damage accumulation is an inevitable consequence of cellular metabolism. When the rate of damage accumulation exceeds the rate at which it is diluted through cell growth and division, a dedicated mitigation strategy is required to prevent population collapse. These mitigation strategies can be coarsely divided into two categories: repair and asymmetrical damage allocation. Repair requires the synthesis of replacement materials for the existing essential machinery or transfer of completely dysfunctional material into the external environment. Asymmetry requires intracellular mass transfer so that, upon division, daughter cells receive different amounts of damage. Perhaps the most common form of somatic damage is the accumulation of misfolded proteins which may be repaired through proteolytic degradation (Dougan et al., 2002) or asymmetrically shuttled and aggregated (Coelho et al., 2013, 2014; Nakaoka & Wakamoto, 2017; Vedel et al., 2016). Other forms of somatic damage are diverse (Nyström, 2003; Starke-Reed & Oliver, 1989; Vigouroux et al., 2020), as are repair and shuttling mechanisms observed among both prokaryotes (Lindner et al., 2008; Schramm et al., 2019; Vaubourgeix et al., 2015) and eukaryotes (Aguilaniu et al., 2003; Coelho et al., 2013; J. Song et al., 2014; Spokoini et al., 2012), including multicellular organisms (Bufalino et al., 2013; Rujano et al., 2006). Investment in dedicated somatic damage mitigation is costly. Under the conceptual framework adopted in this work, repair is explicitly growth-costly whereas asymmetric allocation reduces the fitness of the daughter cell that receives more than half of the somatic damage accumulated by the mother cell. Like the much better understood mechanisms that limit the accumulation of genetic damage, somatic damage mitigation likely plays a central role in the homeostasis of both unicellular organisms and tissues of multicellular ones.

We developed and analyzed a generalizable mathematical model for cell growth, division, and somatic damage mitigation aiming to elucidate the manner in which the optimal strategy depends on four key variables: i) extrinsic, damage-independent mortality, Ii) the efficiency of repair, iii) the rate of somatic damage accumulation, and iv) the model of damage accumulation, ranging from linear to exponential. We modeled the growth of a population of clonal cells within a chemostat until pseudo-equilibrium conditions are met. In order to completely describe the dependence of the damage mitigation strategy on these four factors subject to computational limitations, our framework introduces flexibility with respect to these parameters at the cost of introducing three principal limiting constraints. First, we determine optimal strategy through maximization of an absolute fitness function, the average total number of cells in the population at pseudo-equilibrium. This approach does not differentiate between true equilibria and periodic orbits or limit cycles and does not admit the evaluation of direct competition between pairs of strategies. This also does not admit evaluating trade-offs between stability and adaptability (N. Rochman et al., 2016) which are affected by non-genetic heterogeneity (N. D. Rochman et al., 2018), and it follows, by asymmetry. Second, we assume that asymmetry is not growth costly. By varying repair efficiency, this admits the exploration of conditions where the cost of repair is arbitrarily low, but not conditions where asymmetry is more costly than repair. As described in the Results, this also impacts our observation that intermediate asymmetry is never optimal within our framework; even under conditions where a mixed strategy is optimal, asymmetry is maximal, combined with repair. While consistent with other studies (Clegg et al., 2014), this finding is highly model-dependent (Blitvić & Fernandez, 2020; Evans & Steinsaltz, 2007; Jiseon Min & Amir Ariel, n.d.; Rashidi et al., 2012). These assumptions further do not admit the exploration of conditions where the cost of asymmetric allocation of damage between daughter cells depends on the amount of damage allocated. Third, we introduce extrinsic mortality through dilution of the chemostat which is coupled to nutrient availability.

These constraints did not, however, limit our model agreement with prior efforts. Although not ubiquitously reported (R. Song & Acar, 2019), our model reproduces the central result obtained through both diverse theoretical models (Clegg et al., 2014; Vedel et al., 2016) and experimental inference (Coelho et al., 2013; Vedel et al., 2016), that increasing the rate of somatic damage accumulation tends to increase asymmetric allocation. To our knowledge, this result has been previously obtained only for models in which somatic damage accumulated linearly. Indeed, we observed the same behavior when damage accumulation is linear or approximately linear; however, when damage accumulates strongly nonlinearly, the dynamics are more complex.

Although direct measurements assessing the rate of somatic damage accumulation in single cells remain challenging, model biophysical systems display substantial nonlinearity. In particular, protein misfolding and aggregation follows a “seeding-nucleation” model in which early stages are thermodynamically unfavorable until a stable aggregate is formed at which point growth becomes exponential (Moreno-Gonzalez & Soto, 2011). Similar dynamics are observed over a wide range of model systems including hemoglobin gelation (Ferrone et al., 1980; Saha & Deep, 2014) as well as aggregation of ovalbumin (Roberts, 2007; Weijers et al., 2003) and lactic dehydrogenase (Zettlmeissl et al., 2002). When seed formation, that is, *de novo* misfolding, is the rate limiting step, these dynamics are linear, but when aggregation is rate limiting, they become exponential (Roberts, 2007).

When comparing linear and nonlinear conditions that are equally challenging with respect to the fitness of naïve cells or those adopting the optimal strategy for each condition, we observed that nonlinear conditions are less likely to result in population collapse and that in this case the optimal strategy preventing collapse is more likely to be repair preferred. These effects appear to be a consequence of the fact that, under strongly nonlinear conditions, damage accumulation rate is more robustly modifiable through investment in repair. In addition to mutually exclusive repair and asymmetry preferred mitigation strategies, a mixed optimal strategy including investment in repair and maximal asymmetric allocation is also observed for these conditions of non-linear damage accumulation.

Indeed, this mixed strategy, which generates a subpopulation of cells that inherit no damage upon division through asymmetric allocation and also reduces the rate of damage accumulation through investment in repair, is always optimal when the model of damage accumulation is strongly nonlinear. Under these conditions, increasing the linear rate of damage accumulation (e.g. via an uncorrelated environmental stressor) bifurcates the optimal strategy into either asymmetry preferred when extrinsic mortality is high or repair preferred when extrinsic mortality is low. Fig. 6 provides a graphical summary of these dependencies across all conditions studied.

**Figure 6.**
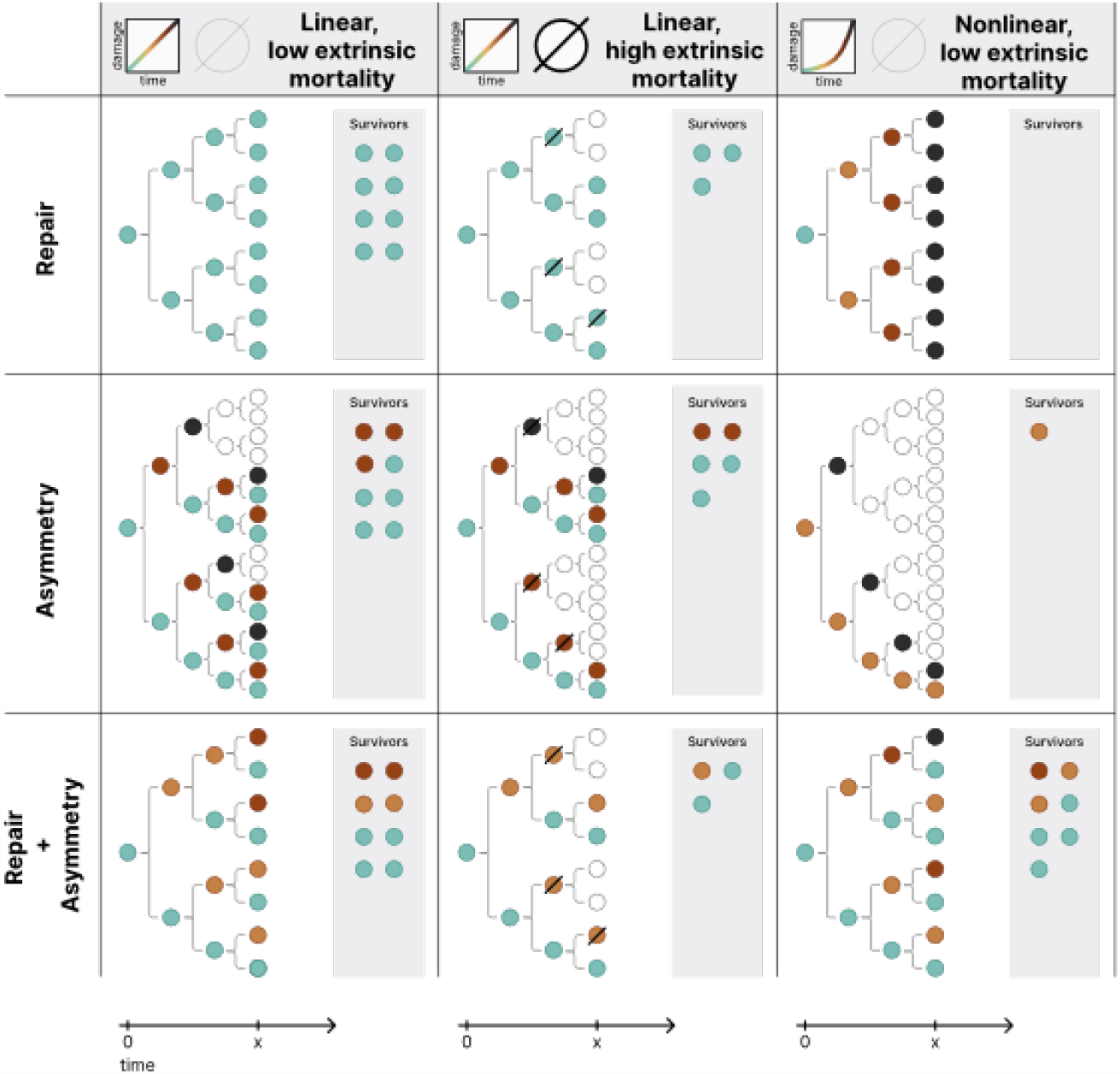
Mitigation strategies under different models of somatic damage accumulation. Rows represent the repair preferred, asymmetry preferred, and mixed strategies. Columns represent linear models of somatic damage accumulation, where the rate of somatic damage accumulation is relatively lower or higher, and a nonlinear model of damage accumulation. Several generations of progeny are illustrated over the same window of observation. Optimal strategy in each conditions is on the diagonal.

The demonstration of the dependence of the somatic damage mitigation strategy on extrinsic mortality is the principal finding of this work. The theory of r/K selection categorizes reproductive strategies with respect to the trade-off between producing the maximum possible number of progeny (r-strategies) and producing (relatively) few high quality offspring (K-strategies). Subject to the maximum number of individuals the environment can support, *K*, and the maximum growth rate of the population, *r*, the population grows according to the rate equation:

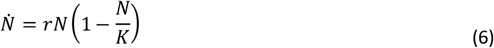

When extrinsic mortality is high, *N* ≪ *K* and r-strategists prevail; the converse is true when extrinsic mortality is low. Unicellular organisms that employ r-strategies tend to have relatively short cell cycle but lower metabolic efficiency, admitting a phase of rapid growth that can be followed by a gradual decline resulting from competition with the more efficient K-strategists (Ho et al., 2017). Specific genomic determinants of substrate preference and efficiency commonly associated with r- and K-strategists have been identified at variable evolutionary depths (Ho et al., 2017). In contrast, relatively little prior work focused on generalizable division strategies which can be realized among diverse species and would r/K selection, as has been explored in greater depth in the related area of cell size regulation (Sauls et al., 2016).

The relationship between growth and efficiency is complex. In microorganisms, the growth rate is strongly cell-density dependent (Ebrahimi et al., 2019; Rosenberg et al., 1977). As observed within our model, this dependence couples *r* and *K*, and can result in co-variance with respect to the changing metabolic strategy rather than a trade-off, but (Marshall et al., 2023). In this work, we demonstrate that somatic damage repair, a K-strategy, is favored when the extrinsic mortality rate is low whereas asymmetric damage allocation, an r-strategy, is favored when this rate is high. Both strategies are employed among the same extant prokaryotes and eukaryotes (Coelho et al., 2013) and were likely accessible to the cellular organisms since very early stages of evolution. We hypothesize that optimization over somatic damage repair and asymmetric allocation played a critical role in establishing repair and asymmetry specialists including the evolution of morphologically asymmetric division. In this sense, the balance of repair and asymmetry in early cellular life seems to represent the prototypical r/K selection regime and perhaps the origins of the first specialized r/K strategists. Furthermore, it appears likely that asymmetric damage allocation, a comparatively simple r-strategy, was the first damage mitigation strategy to evolve in primordial cells that were likely less robust than modern cells, having a high extrinsic mortality rate.

Finally, it is worth briefly discussing how the damage mitigation dynamics shift when somatic damage is externalized (Smakman & Hall, 2022), affecting damage accumulation by cells in addition to inheritance and production. When damage is externalized, the rate of damage accumulation is, essentially, linearized. When damage is not externalized, linearization increases the relative fitness of asymmetry preferred strategies; however, when damage is externalized, the asymmetrically allocated damage can be reintroduced via environmental transfer whereas repaired damage cannot. Within our framework, we observed that the landscape for externalizable damage did not substantially differ from what is discussed in the main text, apparently balancing these opposing influences (Fig. S4). The forms of somatic damage described above may be rarely externalized, viral infection represents a specific, strongly nonlinear form of externalizable somatic damage accumulation. Unicellular organisms evolved a diverse range of viral defense systems to prohibit virion cell entry or destroy viruses intracellularly (Lepikhov et al., 2001; Makarova et al., 2011; Tesson et al., 2022). One mechanism by which cells remove viruses from the population is through programmed cell death (PCD). Asymmetric damage allocation is similar to PCD in that one daughter cell attains a state of reduced fitness (and can die if the damage level is high) relative to competitors that divide symmetrically under the same conditions. In contrast to typical PCD, the other cell attains increased fitness. In this strongly nonlinear regime, the results presented here suggest that mixed damage mitigation strategies should be common, suggesting that PCD is even more widespread in unicellular life forms than presently realized.

## Acknowledgements

The authors are grateful to the Koonin’s laboratory members for valuable suggestions and comments, and to Anastasia Troshina for helping with figures design. Computations were performed using Biowulf HPC cluster of the National Institutes of Health. The authors’ research is funded by the Intramural Research Program of the National Institutes of Health (National Library of Medicine). NDR additionally received intramural support from the City University of New York Graduate School of Public Health and Health Policy (CUNY SPH).

