## Appendices for "Unicellular life balances asymmetric allocation and repair of somatic damage representing the origin of r/K selection"

### Appendix I: ODE solution

We can solve the system of ODEs by setting all the derivatives to 0:

$$0 = \dot{u} = A(1-\frac{r}{E})\phi u - \frac{\rho}{1-\rho}u - Bu$$
$$0 = \dot{\phi} = (1-\phi)B - C\phi u$$
$$0 = \dot{\rho} = D - r + F\rho - A(1-\frac{r}{E})\phi\rho$$

A trivial equilibrium point is immediately apparent: $u = 0, \phi = 1$ (the value of $\rho$ then does not have any meaning). Assuming $u \neq 0$:

$$\frac{\rho}{1-\rho} + B = A(1-\frac{r}{E})\phi \qquad \text{(I1)}$$

$$C\phi u = (1-\phi)B \qquad \text{(I2)}$$

$$A\left(1-\frac{r}{E}\right)\phi\rho = D - r + F\rho \qquad \text{(I3)}$$

Assuming $\rho \neq 0$, from (3) follows $\phi = \frac{D-r+F\rho}{A\left(1-\frac{r}{E}\right)\rho}$. Substituting $\phi$ in (1) with this expression we get

$$\frac{\rho}{1-\rho} + B = A(1-\frac{r}{E})\frac{D-r+F\rho}{A\left(1-\frac{r}{E}\right)\rho}$$

$$(1-B+F)\rho^2 + (B-F+D-r)\rho - (D-r) = 0 \qquad \text{(I4)}$$

Solving (4) yields

$$\rho^* = \frac{-(B-F+D-r)\pm\sqrt{(B-F+D-r)^2+4(D-r)(1-B+F)}}{2(1-B+F)} = \frac{1}{2}(-\frac{B-F+D-r}{1-B+F} \pm \sqrt{\left(\frac{B-F+D-r}{1-B+F}\right)^2 + 4\frac{D-r}{1-B+F}}\ ) \qquad \text{(I5)}$$

which only exists if $D \geq r$ and $B < 1 + F$ or $D < r$ and $B > 1 + F$. We will only consider $B < 1$ (1 is the maximum fraction of a chemostat that can be diluted per unit time) and $D \geq r$ ($r = D$ is the maximum repair that could make sense because for greater $r$, there would be no damage to repair).

The equilibrium value for $\phi$ is then given by substituting $\rho^*$ from (5) to (2):

$$\phi^* = 2\frac{D-r}{A\left(1-\frac{r}{E}\right)}\left(\sqrt{\left(\frac{B-F+D-r}{1-B+F}\right)^2 + 4\frac{D-r}{1-B+F}} - \frac{B-F+D-r}{1-B+F}\right)^{-1} + \frac{F}{A(1-\frac{r}{E})} \qquad \text{(I6)}$$

As we measure the concentration of nutrient in units of its concentration in the source of dilution, it can never exceed 1. That gives us an additional constraint:

$$A\left(1-\frac{r}{E}\right) - F > 2(D-r)\left(\sqrt{\left(\frac{B-F+D-r}{1-B+F}\right)^2 + 4\frac{D-r}{1-B+F}} - \frac{B-F+D-r}{1-B+F}\right)^{-1} \qquad \text{(I7)}$$

When this constraint is not met, the population dies out (steadily increasing $\phi$ implies steadily decreasing $u$). When it is met, the equilibrium population volume size can be expressed from the second equation of the system:

$$u^* = \frac{B(1-\phi)}{C\phi} = \frac{B}{C}\left(2\frac{D-r}{A\left(1-\frac{r}{E}\right)}\left(\sqrt{\left(\frac{B-F+D-r}{1-B+F}\right)^2 + 4\frac{D-r}{1-B+F}} - \frac{B-F+D-r}{1-B+F}\right)^{-1} + \frac{F}{A\left(1-\frac{r}{E}\right)}\right)^{-1} - \frac{B}{C} \quad \text{(I8)}$$

Note that we assumed $\rho \neq 0$ deriving (8). For $D = r = 0$ and $F = 0$, $\rho = 0$ finding equilibrium $\phi$ and $u$ is trivial:

$$\phi^* = \frac{B}{A} \quad \text{(I9)}$$

$$u^* = \frac{B(1-\phi)}{C\phi} = \frac{A-B}{C} \quad \text{(I10)}$$

Population volume size is proportional to the degree to which its growth rate $A$ exceeds death rate $B$ scaled by $C$.

### Appendix II: 1D PDE derivation

A system of ODEs is unsuitable to model asymmetric allocation of somatic damage. That is because when damage is distributed asymmetrically a population subdivides into multiple subpopulations with various damage concentrations instead of being monomorphic like a symmetrically dividing population.

Therefore, instead of modelling two quantities, $n(t)$ and $\rho(t)$, we will now have to model a distribution $n(\rho, t)$ with a PDE. There are three processes that we will have to consider: cell growth and division, cell death and damage accumulation and repair.

Let's write the change of a distribution $n(\rho)$ in a small time $\Delta t$:

$$n(\rho, t+\Delta t) = n\left(\rho - \frac{d\rho}{dt}\Delta t, t\right) + A\phi(t)\left(n\left(\frac{\rho}{1+a}, t\right) + n\left(\frac{\rho}{1-a}, t\right) - n(\rho, t)\right)\Delta t - \left(\frac{\rho}{1-\rho} + B\right)n(\rho, t)\Delta t \quad \text{(II1)}$$

The first term represents the influx to $n(\rho)$ from $n\left(\rho - \frac{d\rho}{dt}\Delta t\right)$ as a result of damage accumulation, the second term represents the growth and division (specifically, influx from the cells that received more damage than their siblings upon division and the cells that received less damage than their siblings upon division and the outflux of the cells that currently have $\rho$ damage and are growing/dividing) and the last term represents death (both from damage, $\frac{\rho}{1-\rho}$, and dilution, $B$). The rate of growth and division is given by $A\phi(t)$ as in the ODE.

Defining $\Delta\rho = -\frac{d\rho}{dt}\Delta \mathrm{t}$ and rearranging:

$$\frac{n(\rho, t+\Delta t)}{\Delta t} = \frac{n(\rho+\Delta\rho, t)}{\Delta t} + A\phi(t)\left(n\left(\frac{\rho}{1+a}, t\right) + n\left(\frac{\rho}{1-a}, t\right) - n(\rho, t)\right) - \left(\frac{\rho}{1-\rho} + B\right)n(\rho, t)$$

$$\frac{n(\rho,t+\Delta t)-n(\rho,t)}{\Delta t}=\frac{n(\rho+\Delta\rho,t)-n(\rho,t)}{\Delta t}+A\phi(t)\left(n\left(\frac{\rho}{1+a},t\right)+n\left(\frac{\rho}{1-a},t\right)-n(\rho,t)\right)-\left(\frac{\rho}{1-\rho}+B\right)n(\rho,t)$$

$$\frac{n(\rho,t+\Delta t)-n(\rho,t)}{\Delta t}=-\frac{d\rho}{dt}\frac{n(\rho+\Delta\rho,t)-n(\rho,t)}{\Delta\rho}+A\phi(t)\left(n\left(\frac{\rho}{1+a},t\right)+n\left(\frac{\rho}{1-a},t\right)-n(\rho,t)\right)-\left(\frac{\rho}{1-\rho}+B\right)n(\rho,t)$$

Taking the limit as $\Delta t\rightarrow 0$:

$$\dot{n}=-(D-r+F\rho-A\phi(t)\rho)n'+A\phi(t)\big(n_{+}+n_{-}-n(\rho,t)\big)-\left(\frac{\rho}{1-\rho}+B\right)n(\rho,t) \tag{II2}$$

Where $\dot{n}\equiv\frac{\partial n(\rho,t)}{\partial t}$ represents the time derivative as above, $n'\equiv\frac{\partial n(\rho,t)}{\partial\rho}$ represents the derivative with respect to damage concentration and $n_{\pm}\equiv n\left(\frac{\rho}{1\pm a},t\right)$.

The equation for $\phi$ changes its form from the ODE only a little, now integrating over $n(\rho)$ to get the population size:

$$\dot{\phi}=B(1-\phi)-C\phi\int_0^1 n(\rho,t)d\rho \tag{II3}$$

### Appendix III: 2D PDE derivation

In the 2D PDE the processes of cell growth and damage accumulation/repair manifest themselves as movement in $(p,q)$ space:

$$n(p,q,t+\Delta t)=n\left(p-\frac{\mathrm{d}p}{\mathrm{d}t}(p)\Delta t,q-\frac{\mathrm{d}q}{\mathrm{d}t}(p,q)\Delta t,t\right)-\left(\frac{\frac{q}{p}}{\frac{2p_0}{Q}-\frac{q}{p}}+B\right)n(p,q)\Delta t \tag{III1}$$

Or, defining $\Delta p=\frac{\mathrm{d}p}{\mathrm{d}t}(p)\Delta t$, $\Delta q=\frac{\mathrm{d}q}{\mathrm{d}t}(p,q)\Delta t$, $\rho=\frac{q}{p}$:

$$n(p,q,t+\Delta t)=n(p-\Delta p,q-\Delta q,t)-\left(\frac{\rho}{\frac{2p_0}{Q}-\rho}+B\right)n(p,q)\Delta t \tag{III2}$$

The first term represents cell growth and damage accumulation/repair, the second term represents cell death both of damage and dilution.

Adding terms to both sides and rearranging:

$$n(p,q,t+\Delta t)-n(p,q-\Delta q,t)-n(p-\Delta p,q,t)+n(p,q,t)+n(p,q,t)-n(p,q,t)$$

$$=n(p-\Delta p,q-\Delta q,t)-n(p,q-\Delta q,t)-n(p-\Delta p,q,t)+n(p,q,t)-\left(\frac{\rho}{\frac{2p_0}{Q}-\rho}+B\right)n(p,q)\Delta t \tag{III3}$$

$$n(p,q,t+\Delta t)-n(p,q,t)+n(p,q,t)-n(p,q-\Delta q,t)+n(p,q,t)-n(p-\Delta p,q,t)$$

$$=n(p-\Delta p,q-\Delta q,t)-n(p,q-\Delta q,t)+n(p,q,t)-n(p-\Delta p,q,t)-\left(\frac{\rho}{\frac{2p_0}{Q}-\rho}+B\right)n(p,q)\Delta t \tag{III4}$$

$$\frac{n(p,q,t+\Delta t)-n(p,q,t)}{\Delta t}+\frac{n(p,q,t)-n(p,q-\Delta q,t)}{\Delta t}+\frac{n(p,q,t)-n(p-\Delta p,q,t)}{\Delta t}$$

$$= \frac{n(p-\Delta p,q-\Delta q,t)-n(p,q-\Delta q,t)}{\Delta t} + \frac{n(p,q,t)-n(p-\Delta p,q,t)}{\Delta t} - \left(\frac{\rho}{\frac{2p_0}{Q}-\rho} + B\right) n(p,q) \quad \text{(III5)}$$

$$\frac{n(p,q,t+\Delta t)-n(p,q,t)}{\Delta t} + \frac{n(p,q,t)-n(p,q-\Delta q,t)}{\Delta q}\frac{dq}{dt}(p,q) + \frac{n(p,q,t)-n(p-\Delta p,q,t)}{\Delta p}\frac{dp}{dt}(p) =$$
$$\frac{n(p-\Delta p,q-\Delta q,t)-n(p,q-\Delta q,t)}{\Delta p}\frac{dp}{dt}(p) + \frac{n(p,q,t)-n(p-\Delta p,q,t)}{\Delta p}\frac{dp}{dt}(p) - \left(\frac{\rho}{\frac{2p_0}{Q}-\rho} + B\right) n(p,q) \quad \text{(III6)}$$

Taking the limit as $\Delta t \to 0, \Delta p \to 0, \Delta q \to 0$:

$$\frac{\partial n(p,q,t)}{\partial t} + \frac{\partial n(p,q,t)}{\partial q}\frac{dq}{dt}(p,q) + \frac{\partial n(p,q,t)}{\partial p}\frac{dp}{dt}(p) = \frac{\partial n(p,q,t)}{\partial p}\frac{dp}{dt}(p) + \frac{\partial n(p,q,t)}{\partial p}\frac{dp}{dt}(p) - \left(\frac{\rho}{\frac{2p_0}{Q}-\rho} + B\right) n(p,q) \quad \text{(II7)}$$

With $\frac{\partial n}{\partial q} = (D-r)p + Fq$, $\frac{\partial \mathrm{n}}{\partial p} = A\left(1 - \frac{r}{E}\right)\phi p$ we get the final form of the 2D PDE:

$$\frac{\partial n}{\partial t} + (D-r)p + Fq\frac{dq}{dt}(p,q) = ((D-r)p + Fq)\frac{dp}{dt}(p) - \left(\frac{\rho}{\frac{2p_0}{Q}-\rho} + B\right) n(p,q) \quad \text{(III8)}$$

### Appendix V: The uniqueness of the non-trivial equilibrium state

Suppose there is more than 1 fixed point with at least two corresponding values for $\phi$: $\phi_1$and $\phi_2$. Without loss of generality consider $\phi_2 > \phi_1$.

We can replace the integral in eq. 3B with total volume, $V$ to get the relationship between $\phi$ and $V$:

$$\frac{\mathrm{d}\phi}{\mathrm{d}t} = B(1-\phi) - C\phi V \quad \text{(V1)}$$

At a fixed point $\phi = \frac{B}{B+CV}$ or $V = \frac{B}{C\phi} - \frac{B}{C}$, therefore the volume monotonically decreases with respect to $\phi$. It follows that $V_2 < V_1$.

By conservation of mass in the chemostat at steady state, the sum of diluted dead mass and diluted living mass is constant. Increasing $\phi$ increases the probability a cell will gain volume before gaining damage for all cells, therefore the ratio of the rates at which living volume / dead volume are produced monotonically increases with respect to $\phi$. Therefore living volume, $V$, increases monotonically with respect to $\phi$, and so $V_2 > V_1$.

By contradiction, $V_2 = V_1$ and $\phi_2 = \phi_1$. There may exist at most one non-trivial fixed point.

### Appendix VI: Details of the numerical scheme for finding an equilibrium state

To determine if asymmetry or repair is preferred in certain conditions defined by fixed values of $(A, B, C, D, E, F)$, we computationally solved the master equation system to obtain the equilibrium population size $\bar{n}$ for multiple combinations of $(a, r)$. This gave us an approximation for a fitness landscape in the form of $\bar{n}(a, r)$. We chose the values for $a$ in a uniform grid from 0 to 1 and values for $r$ from a uniform grid from 0 to $\min(\max(D, F), E)$, then continuing up with a logarithmically increasing step until the population died off or all the damage was repaired.

The simulations were initialized with random starting conditions: population size exponentially distributed with mean $10^5$; nutrient concentration uniformly distributed between 0 and 1; 2D gaussian distribution for $n(p,q)$ with means and variances for $p, q$ uniformly distributed between their minimum and maximum values. We claimed convergence after one of the three criteria was met. If the population size did not change for $t = 1000 * \Delta t_{max}$ where $\Delta t_{max}$ is maximum $\Delta t$ used during the simulation, we used this value as equilibrium estimate. If the population size oscillated with decreasing amplitude around some value that could be consistently predicted based on the newly emerging peaks, we used that value as equilibrium estimate. Finally, if the population size oscillated with non-decreasing and non-increasing amplitude around some value, we claimed cyclic behavior and used geometric mean of the two last peaks as convergence estimate.
