## Supplementary Figures for "Unicellular life balances asymmetric allocation and repair of somatic damage representing the origin of r/K selection"

### Figure S1

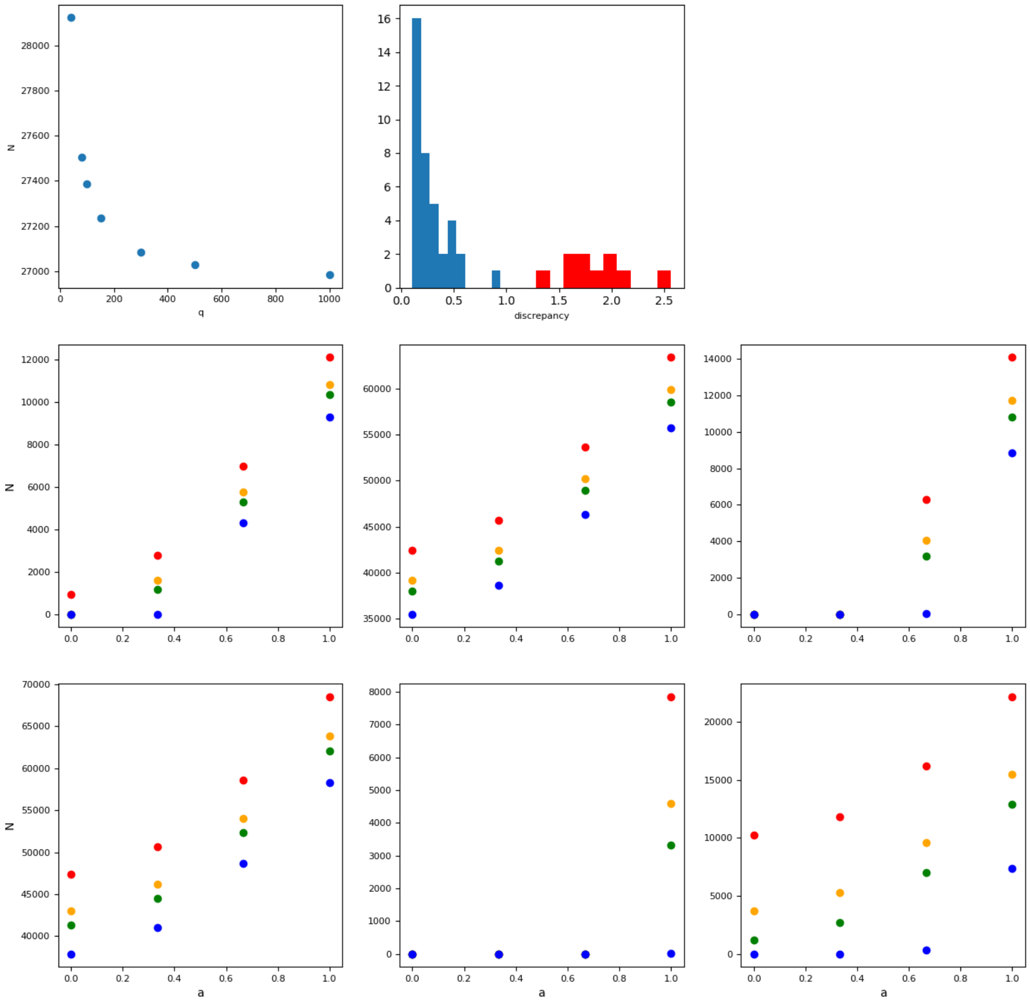


The results are independent of chosen $p$ and $Q$.

The top left panel shows an example of a dependency of equilibrium population size on $Q$. The population size converges as $Q \to \infty$ and at the chosen value $Q = 1000$ the approximate convergence is achieved.

The distribution in the top right corner is the discrepancy between different values of $p \in \{40,\ 80,\ 100,\ 200\}$ calculated as $\frac{\max(N(B_0,D_0,E_0,F_0,a_0,r_0,p)) - \min(N(B_0,D_0,E_0 F_0,a_0,r_0,p))}{\mathrm{mean}(N(B_0,D_0,E_0,F_0,a_0,r_0,p))}$. Most of points (blue distribution) have low discrepancy indicating minimal dependency of the equilibrium population size on $p$. Some of the points (red distribution) show high discrepancy. The fitness landscape slices for r=0 for such cases are shown in the 6 panels at the bottom illustrating that despite of high discrepancy, shape of the landscapes is independent of $p$ (represented by color: 40 – red, 80 – orange, 100 – green, 200 – blue).

### Figure S2

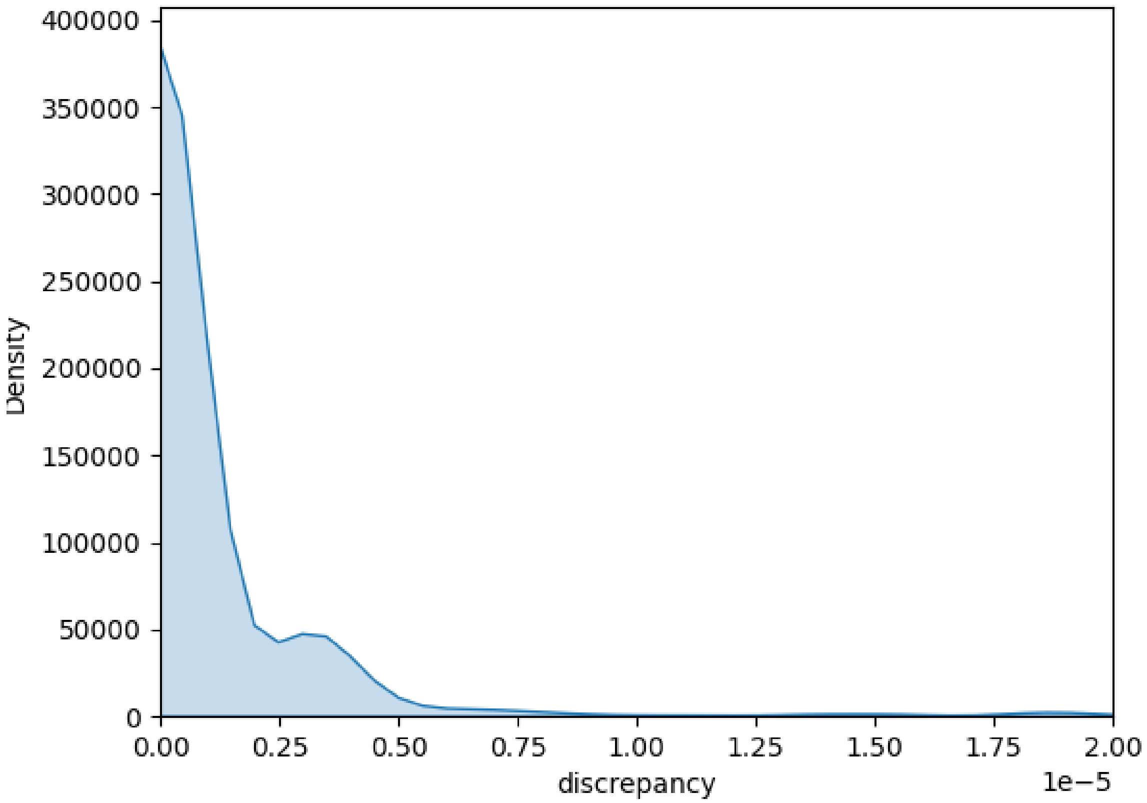


Discrepancy between different values of $C \in \{1\mathrm{e}-5, 1\mathrm{e}-6, 1\mathrm{e}-7, 1\mathrm{e}-8\}$ calculated as $\frac{\max(N\_norm(B_0,D_0,E_0,F_0,a_0,r_0,C)) - \min(N\_norm(B_0,D_0,E_0F_0,a_0,r_0,C))}{\mathrm{mean}(N\_norm(B_0,D_0,E_0,F_0,a_0,r_0,C))}$, where $N\_norm$ is the population size normalized by the population size of a population in the same conditions $B, D, E, F$ adopting a strategy with complete asymmetry and no repair, $a = 1, r = 0$. Comparing the fitness landscapes normalized this way allows to focus on the shape of the landscape rather than its height. The figure illustrates that the discrepancy between landscape shapes was minimal for different values of $C$.

### Figure S3

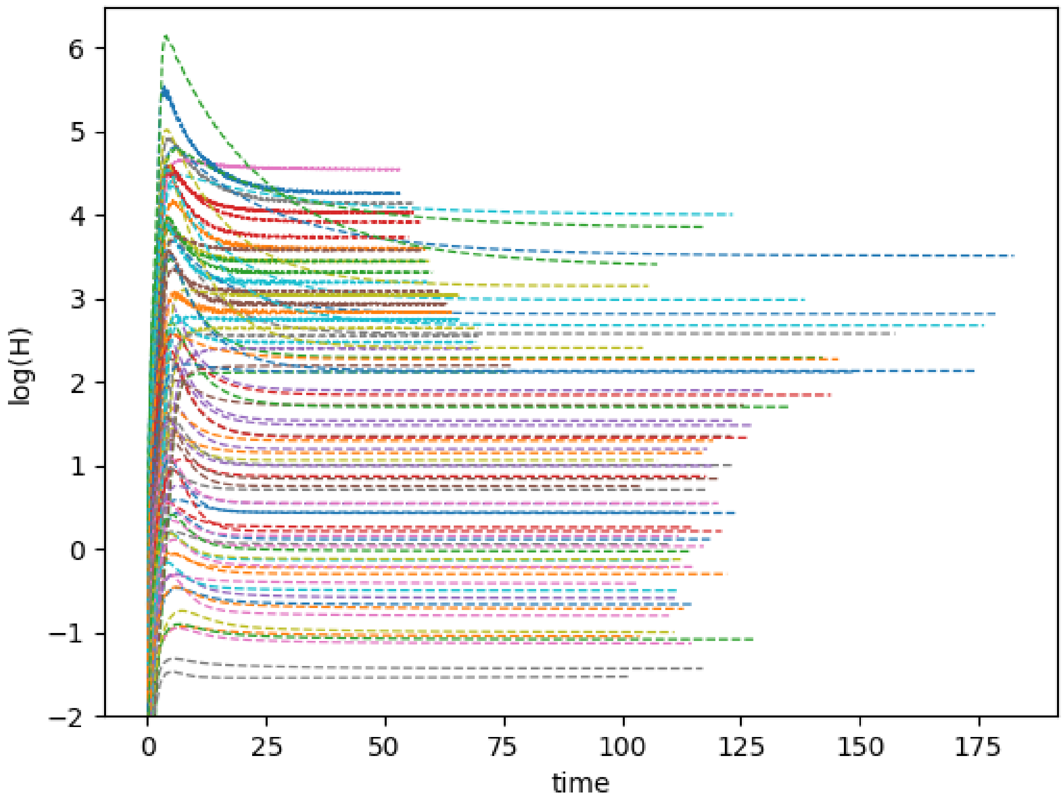


Harshness convergence plots. The figure illustrates that the harshness measures converged for all the conditions studied.

### Figure S4

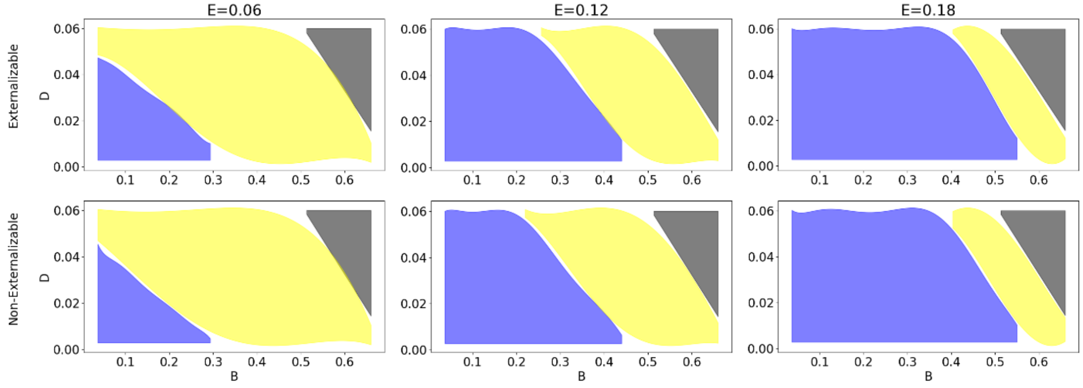


Making damage externalizable does not impact the occupancy of the parameter space by repair/asymmetry strategies.

The option for the damage to be externalized upon cell death with later reabsorption by other cells was incorporated in our 2D master equation model by adding a new variable $\psi$ , number of damage particles in the chemostat. It decreases as they damage is flushed away and consumed by cells, but increases as cells die and release new damage particles into the medium:

$$\frac{d\psi}{dt} = \sum_{p=p_0}^{2p_0-1} \sum_{q} q \frac{\rho}{1-\rho} n(p,q) - \psi B - C\psi \sum_{p=p_0}^{2p_0-1} \sum_{q} pn(p,q)$$

Here we use the same constant $C$, representing the rate of nutrient consumption, assuming that the rate of damage consumption is guided by the same principles.

The single change in the master equation introduced by this is the linear rate of accumulation: instead of being constant it now depends on damage concentration in the chemostat and the rate of its consumption $C\psi$:

$$\frac{dn(p,q)}{dt} = A\left(1-\frac{r}{E}\right)\phi\big((p-1)n(p-1,q) - pn(p,q)\big) - (\frac{\rho(p_0,q)}{1-\rho(p_0,q)} + B)n(p,q)$$

$$+ C\psi p\big(n(p,q-1) - n(p,q)\big) + F\big((q-1)n(p,q-1) - qn(p,q)\big)$$

$$+ rp\big(n(p,q+1) - qn(p,q)\big)$$

### Figure S5

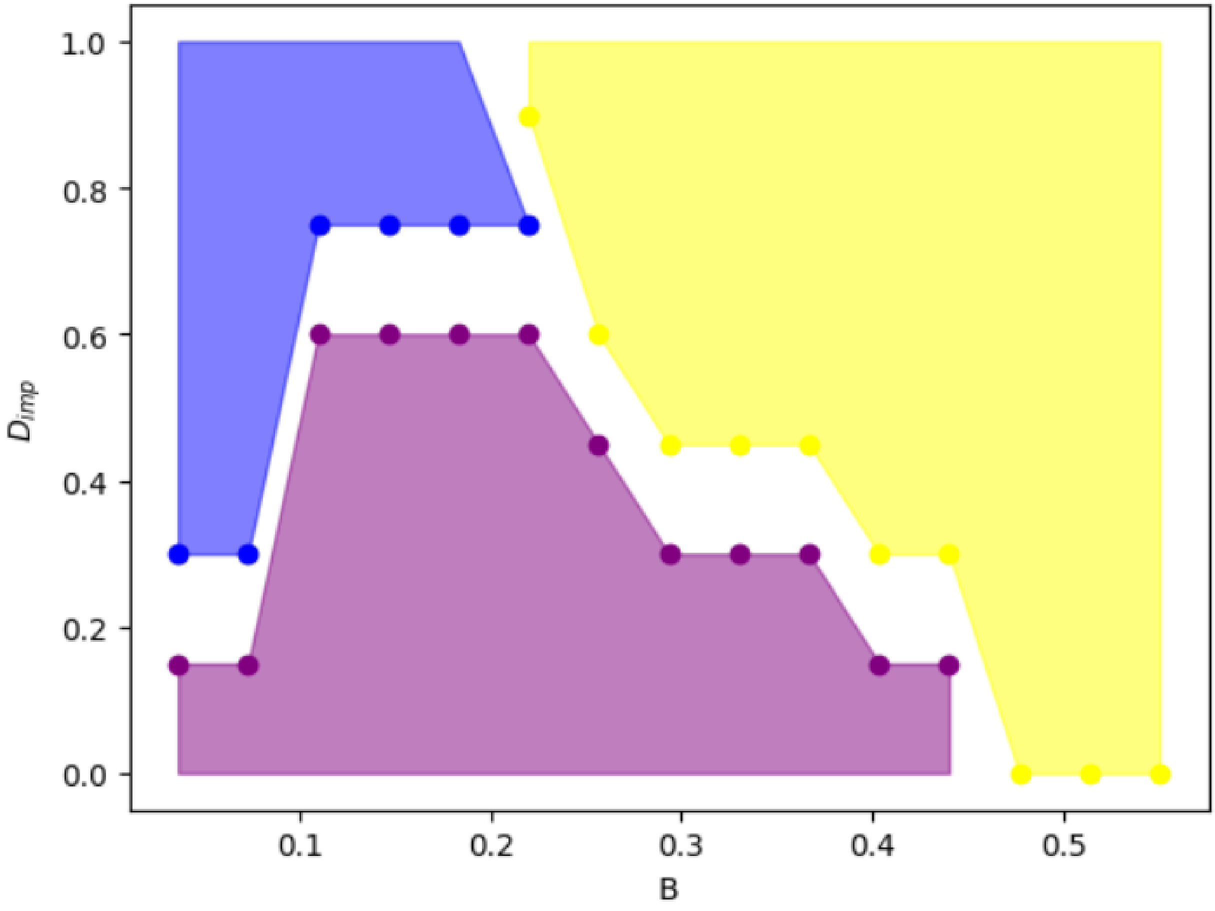


Raw data for Figure 5.
